# Sequential restoring O_2_ then unloading CO_2_ is beneficial against reperfusion injury: role of CO_2_ in metabolism

**DOI:** 10.1101/2023.01.26.525692

**Authors:** Nan Liu, Lv Wang, Ying Xing, Chen Wang

## Abstract

CO_2_ is one of main byproducts during mitochondrial oxidation. Under the acute occlusion of coronary artery situation, the intra-tissue pCO_2_ of heart could be extremely high. This CO_2_ accumulation will be acutely unloaded and discharged by blood reperfusion. However, the effect of this intra-tissue CO_2_ accumulation then unloading process on cardiac ischemic reperfusion injury has not been well investigated yet. In the present study, we show that the perfusion with a high level of pCO_2_ and normal pO_2_ in the initial 30min followed by a 30min normal pCO_2_ and normal pO_2_ is better than the perfusion with 1h normal pCO_2_ and normal pO_2_ simultaneously during the reperfusion after a 45min global ischemia in isolated rat hearts. To observe the effect of high pCO_2_ on cellular metabolism, we exposed C2C12 cells under about 370mmHg pCO_2_ to observe the mitochondrial substrate switch and TCA cycle flux change, by using ^13^C tracers. We show that a short time exposure to the extremely high level of pCO_2_ is not completely destructive for cellular metabolism but has specific effects. The high pCO_2_ inhibits pyruvate transport into mitochondria and the next oxidation, switching to more reliance on fatty acid oxidation and enhancing the glutamine oxidation to maintain the TCA cycle. Intriguingly, the high pCO_2_ significantly activates the reductive carboxylation from glutamine, fixation of mitochondrial excessive CO_2_. The mechanism under the beneficial effect of the high-then-low CO_2_ sequential reperfusion strategy is discussed further.

## Introduction

Glucose and fatty acids serve as the primary substrates that provide acetyl-CoA to the tricarboxylic acid (TCA) cycle in mitochondria. For each mole of O_2_ consumed, there is about 50% higher ATP produced from the complete oxidation of glucose than from fatty acids. Whereas, fatty acids produce significantly more ATP than the same weight of glucose, about 2.4 times, at a greater cost of O_2_. Hence, there is an advantage in utilizing fatty acids for fuel when oxygen is abundant and fuel source is scarce. The reverse, however, would be better using glucose for oxidation when fuel source is plentiful and oxygen is scarce. The ability to efficiently adapt metabolism with specific substrates utilization, according to O_2_ level or substrate availability and energy requirement, is known as metabolic flexibility.

On the other hand, CO_2_ is one of the main products of mitochondrial oxidative metabolism. CO_2_ removal is at least as important as the delivery of O_2_ for cell metabolism and cellular function^1 2^. Normal physiological pCO_2_ is ~40mmHg in artery and ~50mmHg in vein. However, pCO_2_ from skeletal muscle is at least above 120mmHg during heavy exercise^1^. In some pathophysiological conditions such as coronary artery occlusion, an immediate increase of tissue pCO_2_ in myocardium from the baseline 60mmHg to a maximal value of >400mmHg was observed^3 4^. It has been also noted that tissue CO_2_ accumulation is more relative to the myocardial ischemic injury and the intramyocardial ST segment elevation than the O_2_ shortage^5 6^, and with a high regional sensitivity and specifity^7^. The coin of ischemia has two sides, one is hypoxia and the other is hyper CO_2_. Generally, we understand the ischemic reperfusion injury as a hypoxia-reoxygenation process. Whereas, the ischemic CO_2_ accumulation provides a perspective to understand the ischemic reperfusion process, as the “hyper then unloading of CO_2_” process. However, the effect of this intra-tissue CO_2_ accumulation then unloading process on cardiac ischemic reperfusion injury has not been well investigated yet.

What happens if we do not simultaneously restore pO2 and pCO2 to normal levels during reperfusion? We hypothesized that a sequential recovery of pO_2_ then unloading of pCO_2_ might be beneficial for prevention against ischemic reperfusion injury.

## Results

### Sequential reperfusion strategy is beneficial for cardiac function and against reperfusion injury

To test our hypothesis, we used the isolated rat heart model and the global ischemic reperfusion model as before described^8 9^, with Krebs-Henseleit (K-H) medium containing (mmol/ L) 120 NaCl, 25 NaHCO_3_, 4.7 KCl, 1.2 KH_2_PO_4_, 1.2 MgSO_4_, 1.25 CaCl_2_, and 11 glucose (37 °C, pH 7.4), 45min global ischemia followed by 1h reperfusion. We performed the perfusion procedure in a humidified chamber. The atmosphere of the chamber was controlled with a ProCO_2_ carbon dioxide controller (BioSpherix Ltd.). The buffering capacity of the culture medium was modified by changing its initial pH with Tris-MOPS solution to obtain a pH of ~7.4 at the different CO_2_ levels as described previously^10^. We randomized rat hearts into 2 groups, the regular reperfusion group (Reg) and the sequential reperfusion group (Seq). In the Reg group, the reperfusion was directly gassed under the low CO_2_ level (5% CO_2_; pCO_2_: 35–45 mmHg, pH: 7.35-7.45, 50% O_2_/45% N_2_) for 1h. In the Seq group, the reperfusion was first gassed under high CO_2_ level (45% CO_2_; pCO_2_: 360-370 mmHg, pH: 7.35–7.45, 50% O_2_/5% N_2_) for 30min then switched to the low CO_2_ level for the next 30min. After the reperfusion, LDH release was significantly reduced in the Seq group (Fig.1A). Cardiac function was also protected in the Seq group, by measuring the LV systolic pressure (Fig.1B) and LV diastolic pressure (Fig.1C) after 1h reperfusion. The oxidative stress was alleviated significantly, by measuring MDA level (Malondialdehyde, a type of production from lipid peroxidation), or superoxide level (Fig.1D). The level of Glutathione (GSH), the major cellular antioxidant, was significantly higher in the Seq group after 1h of reperfusion (Fig.1E). The cytosolic lactate/pyruvate (L/P) ratio was used as surrogate assay for NADH/NAD+ ratio, a redox indicator. We also found that the L/P ratio was also attenuated in the Seq group (Fig.1F).

**Fig.1.**
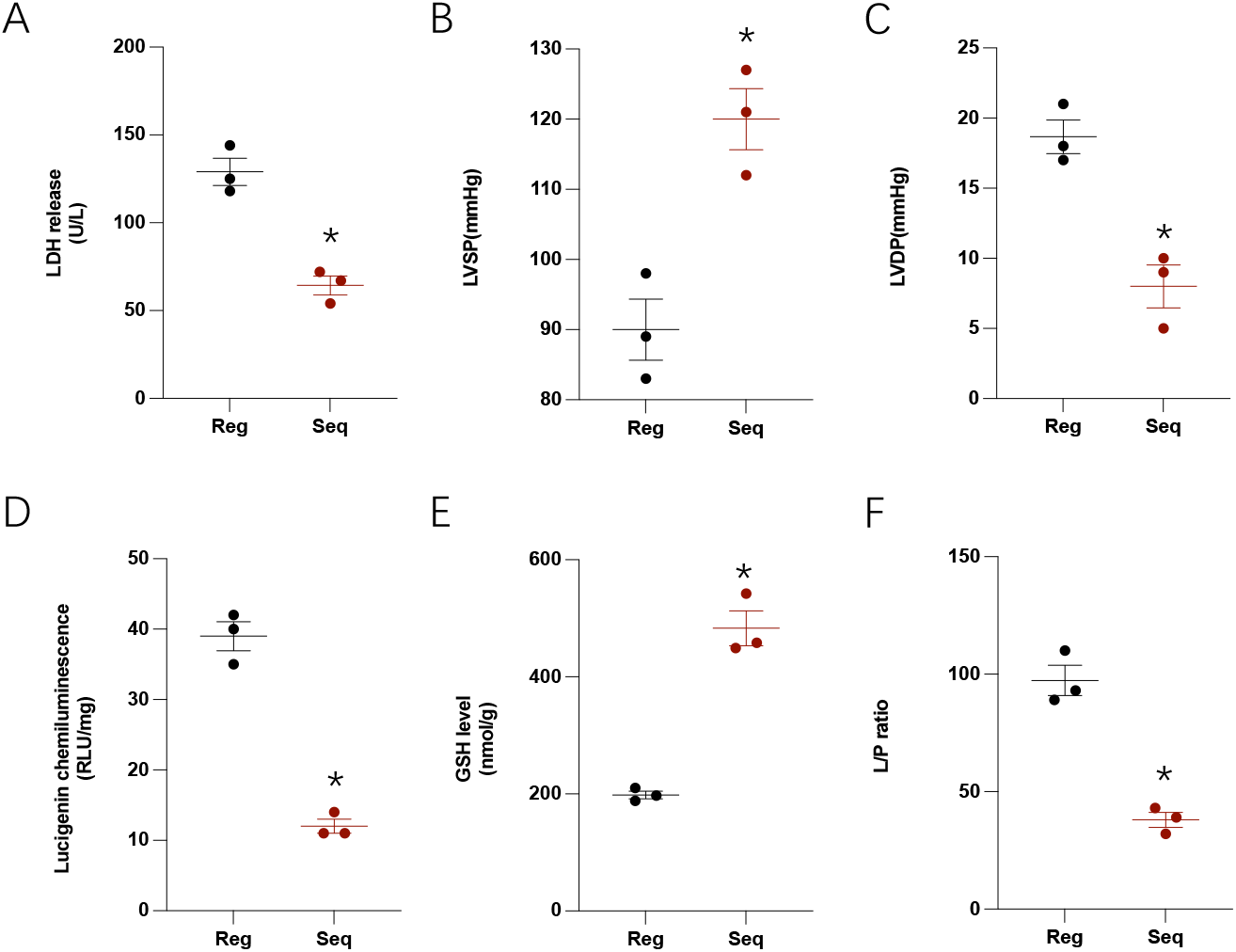
A), sequential recovery of O2 then CO2 level reduced the ischemic reperfusion injury, LDH was collected during the 1h reperfusion. B), left ventricular systolic pressure and C), left ventricular end-diastolic pressure were measured after the 1h reperfusion. D), E) and F) were measured for assessment of oxidative stress. Data are mean±SEM, *p<0.05, by ANOVA with Dunnett’s post hoc test, n=3 for each group.

### High CO_2_ on cellular O_2_ consumption

To investigate how metabolism is affected in response to different levels of CO_2_, we exposed C2C12 myoblasts under Low CO_2_ (5% CO_2_+45% N_2_+50% O_2_) or under High CO_2_ (45% CO_2_+ 5% N_2_+50% O_2_), under room air pressure. The pCO_2_ is ~40mmHg in Low CO_2_ and ~360-370mmHg in High CO_2_. Animal could not live under this extremely high level of CO_2_ but individual organs and local tissues could face this situation, in cardiac muscle^3 4^ or in skeletal muscle^1^. Intriguingly, baseline respiration maintains under High CO_2_, and the ATP-linked respirations were not different between Low and High CO_2_ cells. However, the uncoupler stimulated maximal respiration with all substrates available was significantly decreased under High CO_2_ condition (Fig.2A). These results suggest that the extremely High CO_2_ is not fully destructive for cellular metabolism or halts cellular respiration, but it has a specific effect.

**Fig.2.**
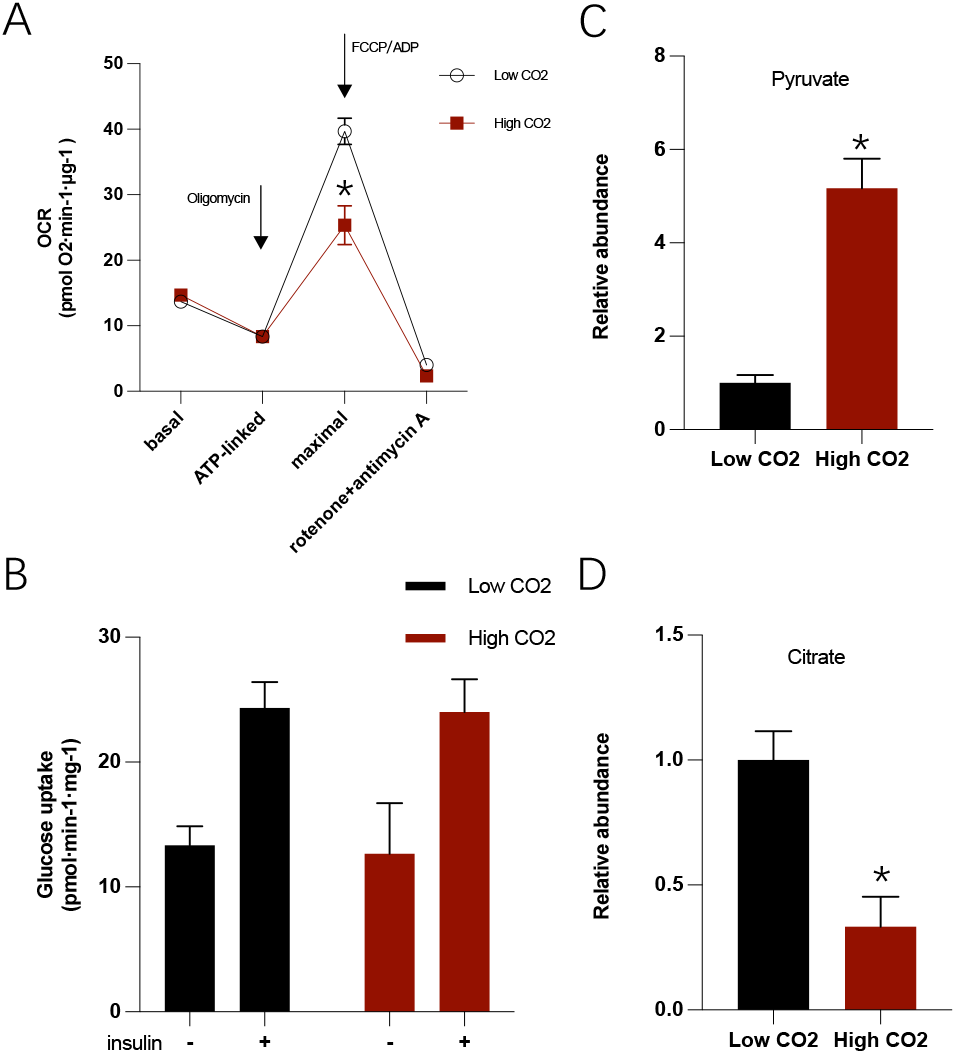
High CO2 effect on general metabolic state of C2C12 cells. A), O2 consumption rates (OCR). Oligomycin sensitive O2 consumption is calculated as ATP-linked OCR. FCCP and ADP act as uncouplers to stimulate maximal OCR. Rotenone and antimycin A act as mitochondrial respiration inhibitors. 4 biological replicates (with a minimum of five technical replicates per experiment). B), Glucose uptake was measured in C2C12 cells after 90min exposure to High or Low CO2, using 2-DG uptake measurement. C) and D), targeted metabolomic analysis was performed to assess the intracellular metabolic changes occurring upon different CO2 levels. Relative abundances of intracellular metabolites were normalized to Low CO2. Data presented as mean ± SEM. n=3 for each group or otherwise stated. *p<0.01 by ANOVA with Dunnett’s post hoc test.

### High CO_2_ effect on glucose uptake and intracellular metabolites

Considering the production of CO_2_ per O_2_, respiratory quotient (RQ), the fatty acid oxidation produces a relatively lower RQ than glucose oxidation. In mitochondria, one mole of glucose-derived pyruvate is converted to one mole of acetyl-CoA with releasing one mole of CO_2_ before entry in the TCA cycle for total oxidation. But if the starting molecule is a fatty acid, it goes through the process of β-oxidation to form acetyl-CoA, which does not generate CO_2_ before entry in the TCA cycle. Therefore, while glucose oxidation is a time-saving and rapid process ^11 12^ with less O_2_-consumption, it produces more CO_2_ with higher respiratory quotient (RQ=1). Using fatty acid over glucose as mitochondrial fuel, RQ=0.7, less CO_2_ will be generated by same amount of oxygen consumed. We hypothesized that cells under the extremely high level of CO_2_ exposure would feedback inhibit the CO_2_ productive way, glucose-pyruvate oxidation, and switch to more reliance on fatty acid as fuel, reducing and slowing CO_2_ production in mitochondrial.

To specifically investigate the effect of High CO_2_ on cellular metabolism, we first measured the glucose uptake using 2-Deoxy-D-Glucose (2-DG). High CO_2_ had no significant effect on basal glucose uptake as well as the insulin-driven glucose uptake in cells measured by 2-DG uptake (Fig. 2B). Next, we measured targeted metabolites in the main metabolic pathways. Total intracellular pyruvate level was increased upon high level of CO_2_ exposure (Fig. 2C), but most of TCA cycle intermediates were not different, except for citrate levels which were decreased under High CO_2_ (Fig. 2D), suggesting a significant decrease of pyruvate-derived acetyl-CoA because majority of citrate comes from pyruvate-derived acetyl-CoA^13^. Aspartate was increased and alanine was decreased, suggesting that amino acid metabolism was affected by High CO_2_.

### High CO_2_ on glucose utilization in cells

While cellular glucose uptake is not affected by High CO_2_, the intracellular glucose metabolism remains unknown. To further analyze intracellular glucose utilization, we cultured C2C12 cells in the presence of [U-^13^C_6_]glucose for 24 hr and observed steady-state isotopic labeling (Fig. 3A). As hypothesized, High CO_2_ decreases the glucose contribution to mitochondrial oxidation as labeling of citrate and all TCA intermediates were significantly decreased in High CO_2_ exposed cells (Fig. 3B and 3C). While the relative abundances of fully labeled (M3) lactate and pyruvate were unchanged in cells under High CO_2_ condition, the extent of alanine labeling from [U-^13^C_6_]glucose was significantly decreased(Fig. 3D).

**Fig.3.**
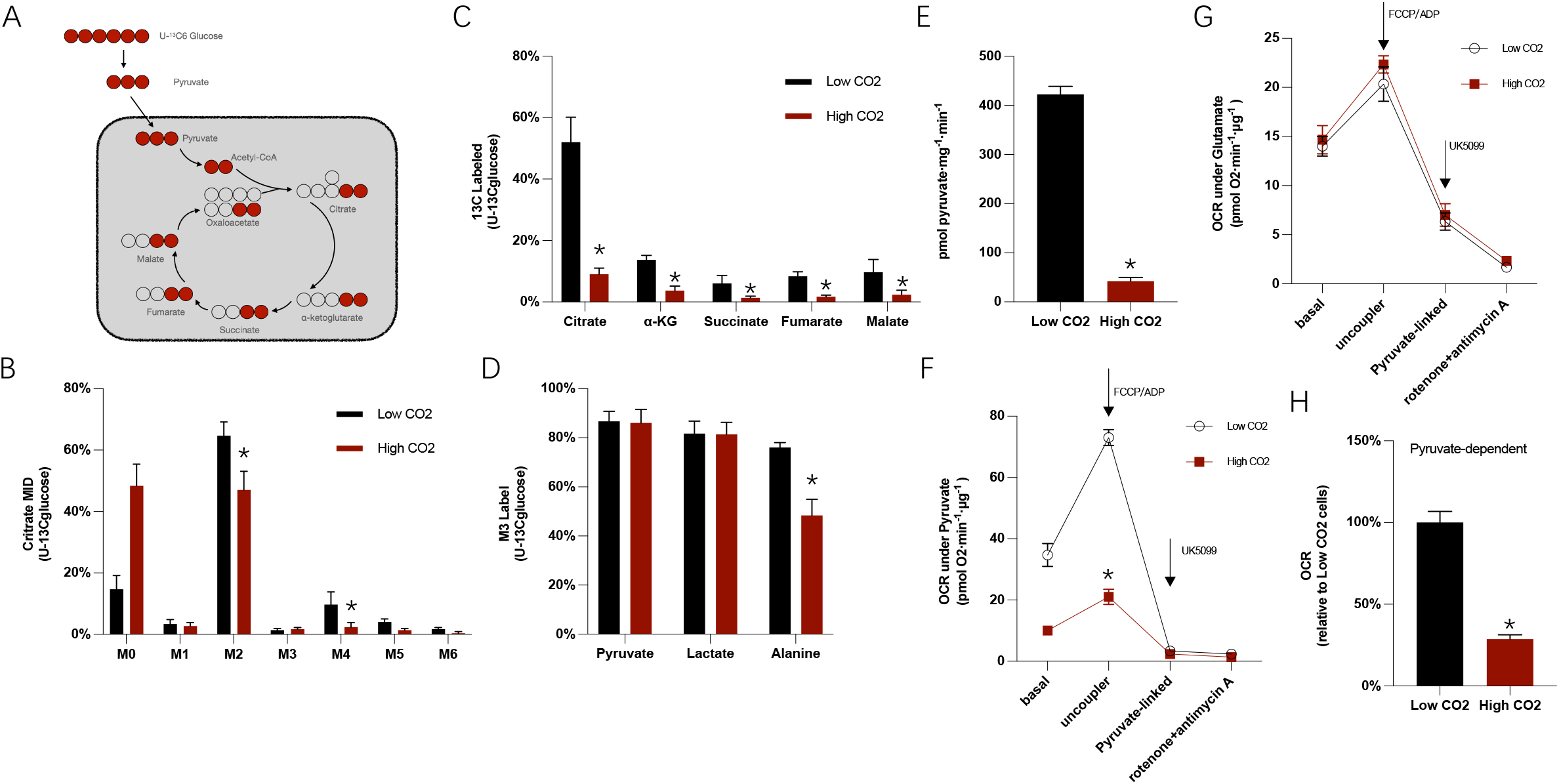
A), Schematic of [U-^13^C_6_]glucose labeling of carbon atoms in TCA cycle intermediates. B), Citrate mass isotopomer distribution (MID) resulting from culture with [U-^13^C_6_]glucose. C), Percentage of 13C-labeled metabolites from [U-^13^C_6_]glucose. D), Percentage of fully labeled lactate, pyruvate, and alanine from [U-^13^C_6_]glucose. E), ^13^C-pyruvate uptake by muscle mitochondria isolated from SD rats, n = 6, two-tailed t-test. F) and G), Pyruvate and glutamate driven respiration by muscle mitochondria isolated from SD rats. Experimental media contained 1 mM malate and 10 mM pyruvate or 10 mM glutamate, 4 biological replicates (with a minimum of five technical replicates per experiment). H), Pyruvatedependent maximal respiration in permeabilized C2C12 cells was significantly compromised. Respiration in permeabilized cells (1 nM XF PMP) was measured in cells offered 5 mM pyruvate, 0.5 mM malate, 2 mM DCA, 2 μg/ml oligomycin, and 400 nM FCCP. n=3 each group or otherwise noted, Data presented as mean ± SEM. *p<0.01 by ANOVA with Dunnett’s post hoc test or otherwise stated.

### High CO_2_ inhibition of mitochondrial pyruvate oxidation

Those results above suggested that High CO_2_ didn’t affect glucose transport and glycolysis but inhibited the next oxidation of pyruvate. To directly assess the pyruvate metabolic change under high level of CO_2_ exposure, we evaluated the mitochondrial pyruvate uptake and pyruvate-driven respiration in the isolated rat skeletal muscle mitochondria. In isolated mitochondria, High CO_2_ inhibited ^13^C-labeled pyruvate uptake (Fig. 3E). The pyruvate-driven, uncoupler stimulated respiration was also markedly compromised (Fig. 3F). In contrast, glutamate-driven, uncoupler stimulated respiration was not affected by high level of CO_2_ exposure (Fig. 3G). In a paralleled permeabilized cell study using C2C12 cells, pyruvate dependent maximal respiration was also significantly compromised under High CO_2_ condition (Fig. 3H).

### High CO_2_ inhibits pyruvate transport

To further investigate the high level of CO_2_ exposure on pyruvate, glutamine and fatty acid metabolism, we quantified the cellular ATP-linked and maximal respiration in the presence of either or all of the inhibitors, UK5099 (MPC inhibitor, inhibiting pyruvate into mitochondria), BPTES (glutaminase inhibitor, inhibiting glutamine oxidation) and etomoxir (CPT1 inhibitor, inhibiting fatty acid oxidation). When cells were treated with UK5099, BPTES or etomoxir, we found high level of CO_2_ exposure had a comparable effect as UK5099 (Fig. 4A, 4B).

**Fig.4.**
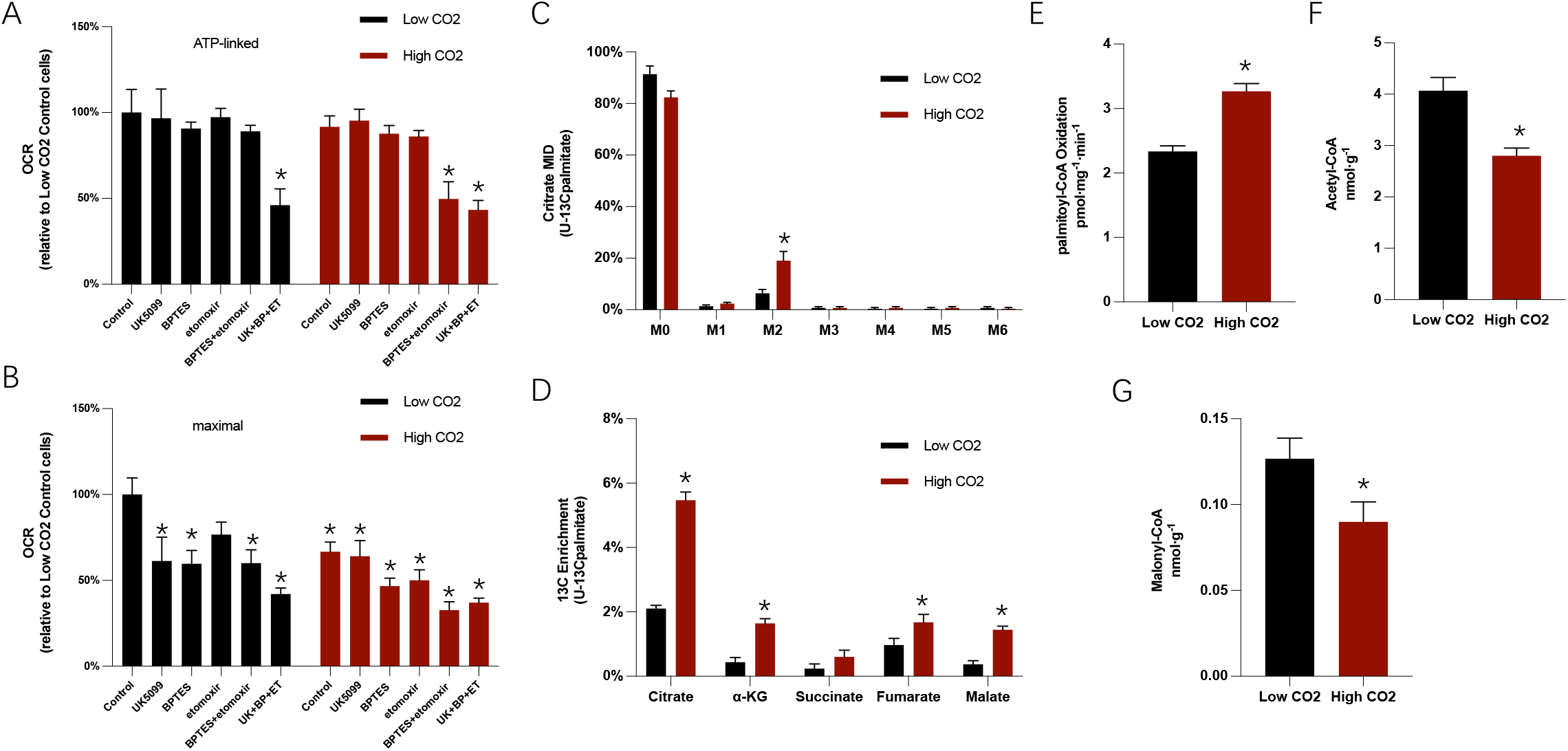
A) and B), ATP-linked and maximal oxygen consumption rate, with or without 20 μM etomoxir, with or without 3 μM BPTES, with or without 10 μM UK5099. Culture medium supplemented with 0.5 mM carnitine. *p <0.01 by ANOVA with Dunnett’ s post. C), Citrate MID resulting from culture with [U-13C16]palmitate conjugated to BSA. *p < 0.01 by two-tailed,equal variance, Student’ s t test. D), Percentage of 13C enrichment resulting from culture with [U-13C16]palmitate. *p < 0.01 by two-tailed,equal variance, Student’ s t test. E), Rate of fatty acid oxidation after addition of palmitoyl-CoA assessed in permeabilized skeletal muscle fibers taken from the quadriceps of SD rats. F) and G),Concentrations of acetyl-CoA and malonyl-CoA in the quadriceps of SD rats. Data are mean ± SEM. n = 4-6 each group. *p <0.01 by ANOVA with Dunnett’ s post.

In Low CO_2_ cells, ATP-linked respirations were only affected when all the three pathways (i.e., pyruvate, glutamine, and fatty acid oxidation) were inhibited (Fig. 4A), while UK5099 was not required in the High CO_2_ cells to decrease ATP-linked respiration (Fig. 4A). On the other hand, maximal respiration was sensitive to each inhibition in Low CO_2_ cells (Fig. 4B). While there was no additional inhibition effect of UK5099 in High CO_2_ cells, the cells were sensitive to BPTES or etomoxir under the High CO_2_ condition (Fig. 4B). These results give us a clear suggest that High CO_2_ inhibits pyruvate transport into mitochondria.

### High CO_2_ increased fatty acid oxidation

As results shown above, the effect of inhibiting the three mitochondrial substrate oxidation pathways shows the mitochondrial metabolism flexibility and suggests that High CO_2_ inhibits the pyruvate transport and drives cell to choose fatty acid and amino acid to meet their oxidation demand. To direct investigate the effect of high level of CO_2_ exposure on fatty acid oxidation, we cultured cells under High or Low CO_2_ levels in the presence of [U-^13^C_16_]palmitate-BSA and observed ^13^C enrichment in TCA intermediates. We observed a significant increase in the relative abundance of M2 citrate from this tracer in High CO_2_ compared to Low CO_2_ cells (Fig. 4C). Increased label incorporation into numerous TCA metabolites downstream of citrate was also detected (Fig. 4D), indicating that High CO_2_ induced a significant increase of fatty acid oxidation in C2C12 myoblasts.

To re-confirm the activating effect of High CO_2_ on fatty acid oxidation, we also directly assessed the rate of fatty acid oxidation after addition of palmitoyl-CoA in permeabilized skeletal muscle fibers taken from the quadriceps of SD rats. It is confirmed that high level of CO_2_ exposure drives an increase in fatty acid oxidation in isolated rat muscles. Rates of fatty acid oxidation were ~25% higher in isolated skeletal muscle under High CO_2_ condition (Fig. 4E). To gain further insight into the effects of High CO_2_ on fatty acid oxidation, we measured concentrations of key allosteric regulators of fatty acid oxidation. Acetyl-CoA produced from pyruvate would be converted to malonyl-CoA, an allosteric inhibitor of carnitine palmitoyltransferase 1 (CPT1, the rate-controlling enzyme of mitochondrial fatty acid oxidation). Indeed, muscle fibers under High CO_2_ condition decreased concentrations of acetyl-CoA (Fig. 4F) and malonyl-CoA (Fig. 4G). To address whether increased fatty acid oxidation might be attributed to changes in the expression of fatty acid oxidation-enzymes, we measured mRNA levels of key fatty acid oxidative-enzymes and found no differences in CPT1, acetyl-CoA carboxylase 2, peroxisome-proliferator-activated receptor α, long-chain acyl-CoA dehydrogenase, and medium-chain acyl-CoA-dehydrogenase between different CO_2_ levels. These findings suggest that decreased inhibition by malonyl-CoA of mitochondrial fatty acid uptake is the mechanism responsible for increased fatty acid oxidation in the skeletal muscle under high CO_2_ exposure.

### High CO_2_ Enhances the contribution of glutamine oxidation to TCA cycle

Besides fatty acid, glutamine oxidation is as well an alternative pathway to compensate glucose derived oxidation in TCA cycle. To analyze mitochondrial glutamine utilization, we cultured C2C12 myoblasts in the presence of [U-^13^C_5_]glutamine to detect changes in glutamine utilization in the context of High CO_2_. The abundance of fully labeled α-ketoglutarate, succinate, fumarate, and malate increased significantly in High CO_2_ cells (Fig. 5A), suggesting that glutamine anaplerosis to TCA cycle increased under High CO_2_ condition. [U-^13^C_5_]Glutamine derived isotopic enrichment in citrate gave us more detail information about the effect of High CO_2_ on TCA cycle metabolic rewiring. In the glutaminolysis pathway, glutamine is converted to α-ketoglutarate entry into TCA cycle and then oxidized by steps to malate (M4) then converted to oxaloacetate (OAA, M4) or to pyruvate (M3). The pyruvate (M3) is further converted by PDH to acetyl-CoA (M2) and then condenses with the OAA (M4) to form citrate (M6) (Fig. 5B)^14^. In our observation, the relative abundance of M6 mass isotopomers was significantly increased in cells under High CO_2_ condition (Fig. 5C). And as we mentioned above, we measured uncoupled respiration in the absence or presence of the glutaminase inhibitor, bis-2-(5-phenylacetamido-1,3,4-thiazol-2-yl)ethyl sulfide (BPTES). Uncoupler-stimulated respiration in High CO_2_ cells was more sensitive to BPTES treatment, suggesting an increased reliance on glutamine oxidation in these cells (Fig. 4B). Collectively, these results provide evidence that High CO_2_ increase glutamine metabolism to maintain oxidation through the TCA cycle.

**Fig.5.**
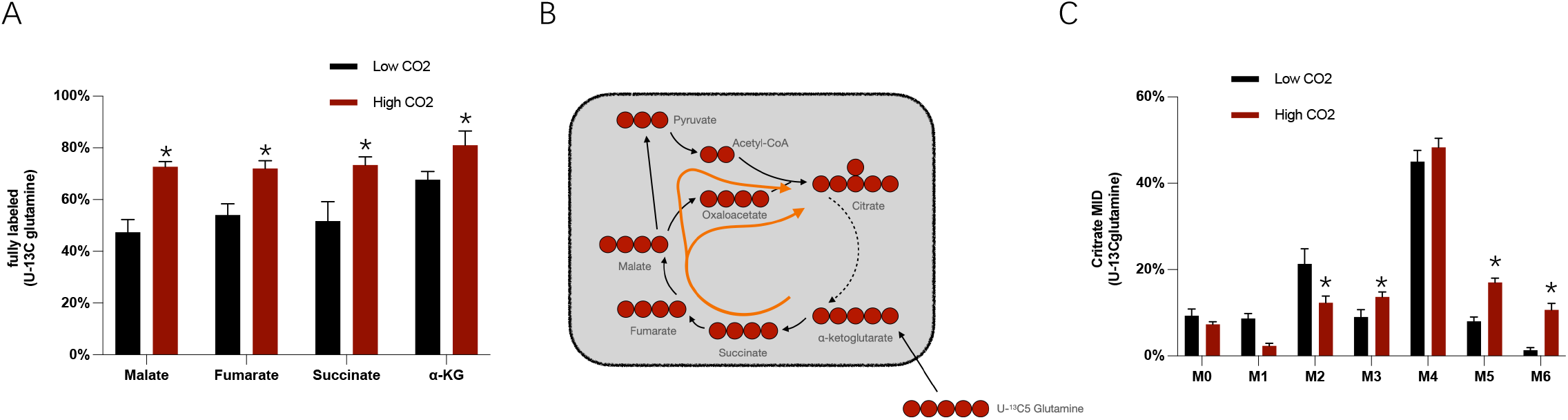
A), Percentage of fully labeled metabolites derived from [U-13C5]glutamine. B), Schematic of [U-13C5]glutamine labeling of carbon atoms in TCA cycle intermediates arising via glutaminoloysis. C),Citrate MIDs resulting from culture with [U-13C5]glutamine. Data are mean ± SEM. n = 4–6 each group. *p <0.01 by ANOVA with Dunnett’ s post.

### High CO_2_ Enhances lipogenesis from reductive carboxylation of glutamine

Intriguingly, the relative abundance of M5 mass isotopomers in citrate was also significantly increased in High CO_2_ cells using [U-^13^C_5_]glutamine tracer (Fig. 5C). The increased M5 citrate abundance is generated via reductive carboxylation by isocitrate dehydrogenase (IDH) enzymes, fixation of one CO_2_ molecule (Fig. 6A)^13^. Then, the citrate could be used for lipogenesis. We quantified isotope enrichment in palmitate and performed isotopomer spectral analysis (ISA) to determine the percent of newly synthesized palmitate after tracer addition and the relative contribution of glutamine to lipogenic acetyl-CoA. Although no significant change in relative palmitate synthesis was observed upon high level of CO_2_ exposure (Fig. 6B), the extent of glutamine conversion to the lipogenic acetyl-CoA pool was significantly increased in High CO_2_ condition (Fig. 6C). Theoretically, glutamine can contribute carbon to citrate and lipogenic acetyl-CoA for fatty acid synthesis via either reductive carboxylation pathway or oxidative glutaminolysis pathway (Fig. 6D). Both the oxidative glutaminolysis pathway and reductive carboxylation pathway were observed in the contribution to the glutamine-derived lipogenesis in Low CO2 cells, but the contribution from reductive carboxylation was significantly increased while the part from oxidative glutaminolysis was not changed significantly in High CO2 exposed cells, using 3-^13^C glutamine and 5-^13^C glutamine as tracers respectively (Fig. 6E).

**Fig.6.**
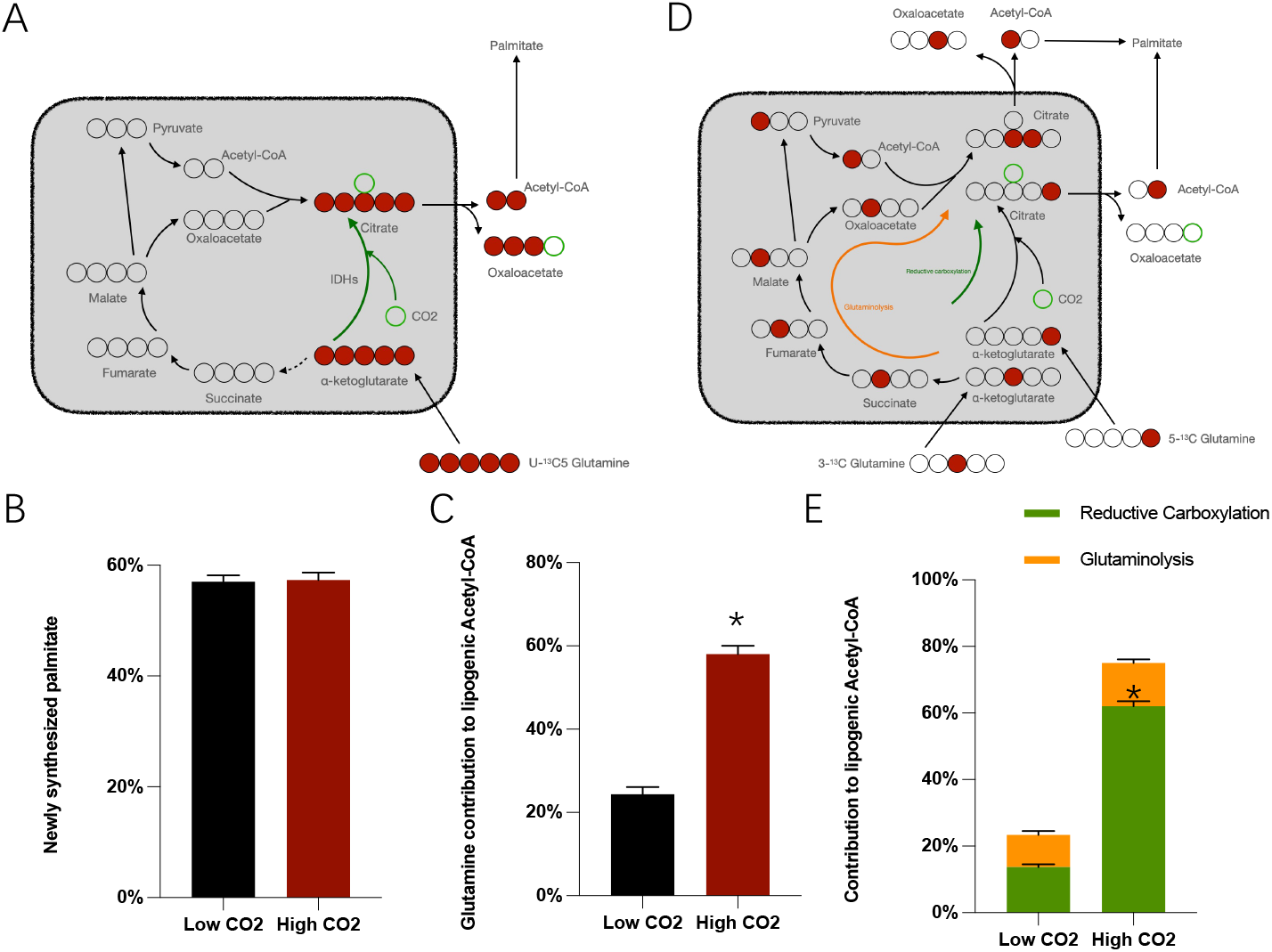
A), Schematic of [U-13C5]glutamine labeling of carbon atoms in TCA cycle intermediates arising via reductive carboxylation and fixation one CO2 molecule. B), Percentage of newly synthesized palmitate as determined by isotopomer spectral analysis. C), Contribution of [U-13C5]glutamine to lipogenic Acetyl-CoA as determined by isotopomer spectral analysis. D), Schematic of [U-13C5]glutamine labeling of carbon atoms in TCA cycle intermediates arising via reductive carboxylation and fixation one CO2 molecule. D), Schematic of contribution of glutamine to lipogenic Acetyl-CoA via glutaminolysis (using [3-13C]glutamine) or via reductive carboxylation (using [5-13C]glutamine). E), Contribution of glutamine to lipogenic Acetyl-CoA via glutaminolysis (isotopomer spectral analysis using [3-13C]glutamine) and reductive carboxylation (isotopomer spectral analysis using [5-13C]glutamine) under Low or High CO2. Data are 95% confidence intervals.

## Discussion

In this report, we show that the sequential reperfusion strategy, restoring O_2_ then unloading CO_2_, is better than the conventional simultaneous reperfusion strategy in the isolated heart model in vitro. CO_2_ is a novel modulator that could regulate the fuel selection and rewire mitochondrial metabolism. High level of CO_2_ exposure could switch mitochondria to the less CO_2_ productive substrates, fatty acids and glutamine, and could also activate a CO_2_ fixation flux, reductive carboxylation. The effects of High CO_2_ on the mitochondrial metabolism occurred within minutes, suggesting a direct, nongenomic effect.

### CO_2_ as a gaseous signal molecule

CO_2_ is produced primarily during aerobic respiration, resulting in higher pCO_2_ levels in mammalian tissues than those in the atmosphere. CO_2_ like other gaseous molecules such as oxygen and nitric oxide, is sensed by cells and contributes to cellular and organismal physiology. Advances in understanding of the biology of high pCO_2_ effects reveal that CO_2_ levels are sensed in cells resulting in specific tissue responses, including the transcriptional responses and cellular and tissue functional responses to elevated pCO_2_ levels^15^.

While the whole-body severe elevation of pCO_2_ is quickly lethal, chronic mild elevation of CO_2_ concentration (hypercapnia) occurs in patients with severe lung diseases. Chronic exposure to that mild elevation of pCO_2_ (<120mmHg) has been investigated. In some observations, mild hypercapnia shows protective effects^16 17 18 19 20^, but other studies get opposite evidences that high pCO_2_ may be detrimental^21 22 23^.

Acute hypercapnia modulates vascular resistance and regulates hemodynamics, with an pH-independent mechanism^24 25 26^. In some earlier studies, dual effects of CO_2_ were observed with both vasoconstriction effect and vasodilation effect on vascular tone ^27 28^. Acute hypercapnia increased vascular resistance in the renal and segmental arteries^29^. pCO_2_ could also regulate cerebral blood flow by dilate or constrict cerebral arteries^30 31^. The human brain generates about 20% of total body CO_2_ production and the effective removal of this CO_2_ relies on the cerebral circulation. When CO_2_ production increases as neuronal activity increases, CO_2_ also acts as a signaling molecule in the neurovascular coupling, regulating local cerebral blood flow in response to increased neuronal CO_2_ production^32^. Furthermore, CO_2_ signaling in regulating cerebral blood flow has a pH-independent mechanism^33 26^. AMPK is one of possible targets of the mild elevation of pCO_2_^23 22^. CO_2_ has comprehensive effects on the heart and skeletal muscle^4 15 34^. High pCO_2_ levels cause skeletal muscle atrophy via an AMPK and FoxOs dependent mechanism^35^ and downregulates skeletal muscle protein anabolism via an AMPK dependent mechanism^23^. Transcutaneous delivery of CO_2_ improves the performance of endurance exercise^36^, improves contractures after spinal cord injury^37^, and accelerate muscle injury repair in rats^38^. These studies suggest that CO_2_ acts as an independent signal molecule playing an important and fundamental role in cellular biology.

### CO_2_ acts on cellular mitochondria and metabolism

The mild elevation of pCO_2_ causes mitochondrial dysfunction and impairs cell proliferation^10^. It was also reported that the elevated pCO_2_ increased skeletal muscle fatty acid oxidation^39^. High pCO_2_ also acts as signaling to improve mitochondrial function with a fuel shift into a substrate-rich, fuel-efficient metabolic mode that allows muscle cells cope with the CO_2_ toxicity^40^. Hypercapnia participate in the skeletal muscle mitochondrial dysfunction during COPD^41^. High pCO_2_ could alter the mitochondrial function, and mitochondria are key players in CO_2_ sensing^19^. Exogenous CO_2_ by transcutaneous delivery improves the performance of endurance exercise and increases the mitochondrial DNA content in skeletal muscle of rats^36^. Hypercapnia also induces mitochondrial dysfunction of mesenchymal stem cells^42^. Acute exposure to hypercapnia induces rapid and substantial reductions in contractile force of the inferior limb muscles. Hypercapnia also decreases ATP production in hyperoxic N12 cells^10^ and in rabbit skeletal muscle^43^. Mask inhalation induced CO_2_ retention in surgery could induce acutely decreased pyruvate oxidation capacity in skeletal muscle mitochondria^44^. These results indicate high pCO_2_ has acute or chronic effect on mitochondrial metabolism.

### High CO_2_ on metabolism rewiring

The previous studies above took aim at the mild level of high pCO_2_ or hypercapnia, which are in the range of clinical values coming from blood samples. The mild level of high pCO_2_ is a chronic and systemic factor throughout the whole body, which is often seen in lung patients. However, the level of pCO_2_ is measured from blood sample, not in local tissue. Indeed, tissue pCO_2_ level could be much higher than it in the venous blood sample, under some pathophysiological situations. After coronary artery occlusion, it was observed an immediate increase of tissue pCO_2_ in myocardium from the baseline 60mmHg to a maximal value of >400mmHg^3 4^. PCO_2_ from skeletal muscle is at least above 120mmHg during exercise^1^, and this value is only got from venous blood but not from the direct tissue measurement. This extremely high level of pCO_2_ (pCO_2_>360mmHg) might play a key role in some pathophysiological process, such as ischemic reperfusion. So, we used the extremely High CO_2_ to observe the direct effect on cell metabolism. While unexpectedly, ATP-linked respiration is unchanged by High CO_2_, we identified specific changes in glucose, amino acid and fatty acid metabolism. Glucose oxidation in the TCA cycle was significantly decreased and pyruvate accumulated in cytosol at the same time. Enhancement of amino acid and fatty acid oxidation were observed upon High CO_2_. Elevated glutamine anaplerosis and oxidation are a consequence of decreased citrate synthase downstream of PDH while the ETC remained active^45 46 47^. We initially hypothesized the fatty acid oxidation might be increased to substitute glucose oxidation under High CO_2_ as the RQ (CO_2_ production) of fatty acid oxidation is lower than that of glucose. That was proved by tracing with [13C]palmitate-BSA in myoblasts and reconfirmed by using palmitoyl-CoA in permeabilized skeletal muscle fibers. The β-oxidation was stimulated, in part, due to decreases in malonyl-CoA, which inhibits CPT1. High CO_2_ effect seems similar with UK5099, suggesting pyruvate transport is inhibited by High CO_2_, whereas PDH activity remains integrity because increased M6 citrate observed as [U-^13^C_5_]glutamine added as tracer.

Intriguingly, we found that High CO_2_ elicited a metabolic state partly similar with hypoxia as well. High CO_2_ enhances reductive carboxylation, which can be observed in the hypoxic metabolic phenotype ^13 14 48 49^. Under the High CO_2_, we found glutamine derived citrate through the reductive carboxylation flux was significantly enhanced than it in Low CO_2_ cells. These citrates come from the glutamine derived α-KG (α-ketoglutarate) with fixation of CO_2_ by IDH (isocitrate dehydrogenase). Then the citrates could be further transported out of the mitochondria, which exhibits the ability of mitochondria to disposal of redundant CO_2_ under some extreme conditions. Significant decrease of alanine level was observed in High CO_2_ cells, which was consistent with a role for mitochondrial pyruvate and mitochondrial alanine aminotransferase (ALT2) in mediating glutamine anaplerosis^50 45^. The accumulation of aspartate in High CO_2_ condition suggests a decreased pyruvate-derived acetyl-CoA and an accumulation of oxaloacetate^45^.

Our results show that extremely High CO_2_ is not a completely inhibitory or destructive factor for cell metabolism, whereas it acts as an acute inhibitor of pyruvate transport and switches the mitochondrial fuel from pyruvate to fatty acids and amino acids to reduce the production of CO_2_, and activates the reductive carboxylation to fix excessive CO_2_ in mitochondria.

### A possible mechanism of CO_2_ in the ischemic reperfusion injury and tumor growth

Understanding the effect of this extremely high level of CO_2_ on cellular metabolism help us to further understand some pathophysiological processes, such as the well-known ischemic reperfusion injury. Generally, we understand the ischemic reperfusion injury as a hypoxia-reoxygenation process. Whereas, our present results provide a new perspective to understand the ischemic reperfusion, as an acute “hyper then unloading of CO_2_” process. As shown in above results, High CO_2_ did not inhibit the cellular respiration and O_2_ consumption but shifted from glucose oxidation to a less CO_2_ production metabolic mode, fatty acid and amino acid oxidation which, however, demanded more O_2_ consumption. So, High CO_2_ could deteriorate the hypoxia. Secondly, High CO_2_ induced pyruvate accumulation in cytosol which would feedback inhibit glycolysis. Whereas, the glycolysis is a protective pathway to generate energy under the hypoxia situation. Therefore, High CO_2_ diminishes the cellular autonomously protective mechanism under hypoxia. Furthermore, to incorporate more CO_2_, carboxylation of α-ketoglutarate to form citrate is enhanced and then citrate diffuses out of mitochondria to cytosol instead of CO_2_, whereas, the acidity of citrate is much higher than carbonate. Accordingly, High CO_2_ worsens the acidosis more severe than CO_2_ per se and deteriorates the glycolysis induced lactate acidosis. On the other hand, during the reperfusion process, CO_2_ level is sharply discharged by blood flow, the inhibitory effect of pyruvate transport is eliminated acutely and O_2_ supply is restored, which might induce an intense oxidation of pyruvate in mitochondria, eliciting too much ROS production and oxidative stress. These would all induce additional injury in the ischemic reperfusion. Base on this point, we can explain the beneficial effects of the sequential reperfusion strategy on the isolated rat hearts. Additionally, it is well known that reductive carboxylation supports growth in tumour cells with defective mitochondria^49^. As shown in our present study, High CO_2_ could activate the reductive carboxylation flux. So, we further suppose that High CO_2_ might also play a role in the tumor growth.

## Methods

### Animals

All animal study protocols were performed according to the National Institutes of Health Guidelines for the Use of Laboratory Animals and were approved by the Ethics Committee on Animal Use and Care. SD rats were purchased from animal center of our university. Rats were housed in a temperature-controlled environment (22°C) with a 12/12 h light/dark cycle and had access to food and water ad libitum. The experimenters were blind to group assignment and outcome assessment. All procedures were designed and performed to minimize animal suffering while respecting the ARRIVE principles.

### Isolated rat hearts

SD rat hearts were isolated and perfused as described previously^8^.

### Cell culture

C2C12 (a mouse myogenic cell line) myoblasts were proliferated in growth media consisting of DMEM and 20% FBS in 5% CO_2_ and 37°C. When myoblasts were grown to confluence, the media was changed to differentiation media consisting of DMEM and 2% horse serum. The differentiation media was changed every 2 days. After 5–7 days in differentiation media, myotubes with multiple nuclei were formed^51^.

### Mitochondrial isolation

Mitochondria from rat skeletal muscle were isolated as described previously^52^. Briefly, the quadriceps muscles of a SD rat were dissected, and mitochondria were isolated essentially. The tissue was minced on ice in 5-mL volumes of cold IBm1 buffer [67 mM sucrose, 50 mM Tris·HCl, 50 mM KCl, 10 mM EDTA, and 0.2% (wt/ vol) BSA; pH 7.4]. The supernatants were removed and replaced with 5 mL IBm1 + 0.05% (wt/vol) trypsin. After 30 min (on ice), the samples were centrifuged at 200 × g at 4 °C for 3 min, after which the pellets were resuspended in 15 mL of chilled IBm1. The samples were homogenized on ice with seven strokes at 70 rpm in a Teflon/glass Potter-Elvehjem homogenizer and centrifuged at 700 × g for 10 min at 4 °C. The resulting supernatants were centrifuged at 8,000 × g for 10 min at 4 °C. The resulting pellets were washed in 5 volumes of IBm2 buffer (250 mM sucrose, 5 mM EGTA, and 10 mM Tris·HCl; pH 7.4) and recentrifuged at 8,000 × g for 10 min at 4 °C. The mitochondria-enriched pellets were gently resuspended in a minimal amount of IBm2 buffer, and protein concentration was measured using a Bradford assay.

### CO_2_ exposure

Cell lines or isolated mitochondria were exposed under low CO_2_ level (5% CO_2_; pCO_2_: 35–45 mmHg, pH: 7.35-7.45, 50% O_2_/45% N_2_), high CO_2_ level (45% CO_2_; pCO_2_: 360-370 mmHg, pH: 7.35–7.45, 50% O_2_/5% N_2_). The buffering capacity of the culture medium was modified by changing its initial pH with Tris-MOPS solution to obtain a pH of ~7.4 at the different CO_2_ levels as before described^23 10^. The desired CO_2_ and pH levels were achieved by equilibrating the medium overnight in a humidified chamber. The atmosphere of the chamber was controlled with a ProCO_2_ carbon dioxide controller (BioSpherix Ltd.). In this chamber, cells were exposed to the desired pCO_2_ while maintaining 50% O_2_ balanced with N_2_. Before and after CO_2_ exposure, pH, pCO_2_, and oxygen tension levels in the medium were measured using an i-STAT blood gas analyzer (Abbott Point of Care). Experiments were started by replacing the culture medium with the CO_2_-equilibrated medium and incubating it in the chamber for the desired time. In a separate study, we compared the effect of 95% O_2_+5% N_2_ versus 50% O_2_+50% N_2_ on the cell metabolism. There was no difference in metabolism observed so the variation of N_2_ and O_2_ within this range didn’t change the cell metabolic state.

### Cell ^13^C tracing

Phenol red-free DMEM was formulated by replacing the substrate of interest with ^13^C-labeled glucose, glutamine, or pyruvate with other components unlabeled. Cultures were washed with PBS before adding tracer media for 15–30 hr unless otherwise specified. Fatty acid oxidation studies were conducted using [U-^13^C_16_]palmitate noncovalently bound to fatty acid-free BSA. [U-^13^C_16_]palmitate-BSA was added to culture medium at 5% of the final volume (50 mM final concentration) with 1 mM carnitine in medium formulated with FBS that was delipidated using fumed silica (Sigma) according to the manufacturer’s instructions.

### Skeletal mitochondrial pyruvate uptake assay

The pyruvate uptake protocol was based on previously published methodology.

Mitochondria were resuspended to 5.0–9.0 mg/mL in Uptake Buffer (120 mM KCL, 5 mM KH_2_PO_4_, 1 mM EGTA, 5 mM HEPES pH 7.4, 1 mM Rotenone, and 1 mM Antimycin A) and were divided into two equal aliquots treated with 2 mM a-Cyano-4-hydroxycinnamic acid (CHC) or vehicle. 20 mL of treated mitochondria were rapidly mixed with 20 mL of 2X Pyruvate buffer (Uptake Buffer, pH 6.2 with 0.10 mM ^13^C-Pyruvate, gassed with high CO_2_ 50% O_2_/50% CO_2_ or with low CO_2_ 50% O_2_/5% CO_2_/45% N_2_) generating the pH gradient needed to initiate uptake in a high CO_2_ condition or a low CO_2_ condition. After 1 min, 80 mL of Stop buffer (Uptake Buffer pH 6.8 supplemented with 10 mM CHC) was rapidly mixed with the samples to halt uptake and open to normal room air condition. Mitochondria were recovered by passing the solution through dual filter system consisting of a 0.8 mM cellulose filter and a 0.45 mM nitrocellulose filter. Filters were washed twice with 200 mL Wash buffer (Uptake Buffer pH 6.8 supplemented with 2 mM CHC and 10 mM Pyruvate). Excess filter material around the separated and washed mitochondria were removed, and filters containing mitochondria were placed into scintillation vials for quantification. Mitochondria pre-treated with CHC were used as a negative control and counts were subtracted from non-pre-treated mitochondria. Samples were normalized to the amount of mitochondrial protein used.

### Cellular glucose uptake assessment

Glucose uptake was measured as described previously^53^. Differentiated C2C12 cells were washed in serum-free medium and incubated ± insulin (30 nM) under low or high pCO2 levels for 90 min at 37 °C in an incubator. Glucose uptake, quantified using the nonmetabolized radiolabeled analog 2-Deoxy-D-Glucose (2-DG, 10 μM final concentration), was measured in triplicate over 10 min at room temperature. Data were normalized to the protein content in each well. The uptake of labeled L-glucose was used to correct samples for the nonspecific diffusion of tracer.

Fatty acid oxidation measurement in isolated permeabilized rat skeletal muscles Quadriceps skeletal muscle fibers were prepared according to a protocol as previously described^54 55^. Briefly, a subsample of the dissected muscle of ~5 mg wet weight was immediately transferred onto a small Petri dish on ice containing 2 mL of ice-cold organ preservation solution (BIOPS) composed of 2.77 mM CaK_2_ EGTA buffer, 7.23 mM K_2_ EGTA buffer, 0.1 μM free calcium, 20 mM imidazole, 20 mM taurine, 50 mM 2-(N-morpholino) ethanesulfonic acid hydrate (Mes), 0.5 mM DTT, 6.56 mM MgCl_2_·6H_2_O, 5.77 mM ATP, and 15 mM phosphocreatine (pH 7.1). The muscle sample was then gently dissected using forceps with sharp tips. To ensure complete permeabilization the fibers were incubated on a shaker at 4 °C in BIOPS solution containing 50 μg/mL saponin for 30 min. Fibers were washed for 10 min at 4 °C in ice-cold mitochondrial respiration medium 5 [MiR05; 0.5 mM EGTA, 3 mM MgCl_2_, 60 mM K-lactobionate, 20 mM taurine, 10 mM KH_2_PO_4_, 20 mM Hepes, 110 mM sucrose, and 1 g/L BSA essentially fatty acid free, adjusted to pH 7.1^55^] with high pCO_2_ or low pCO_2_ levels, and wet weight of the fibers was measured on a microbalance (Mettler Toledo). Subsequent respiration measurements were performed in MiR05 containing 2 mM carnitine at 37 °C using the high-resolution respiromter Oxygraph-2k (Oroboros). A substrate-uncoupler-inhibitor titration protocol was used to assess fatty acid oxidation after addition of palmitoyl-CoA (50 μM, all concentrations are final) with malate (2 mM) and saturating ADP concentrations (3 mM) present.

### Oxygen consumption measurement in cells

Respiration was measured in adherent monolayers of C2C12 myocytes using a Seahorse XF96 Analyzer. Myoblasts were plated at 1×10^4^ cells/well and grown for 2 days. Cells were assayed in unbuffered DMEM (Sigma) supplemented with 8 mM glucose and 3 mM glutamine. ATP-linked respiration was calculated as the oxygen consumption rate sensitive to 2 μg/ml oligomycin. Maximal respiration was calculated as the difference between uncoupler-stimulated respiration (measured as the highest rate from sequential additions of FCCP and ADP) and nonmitochondrial respiration (measured after addition of 2 μM rotenone and 2 μM antimycin A). Where indicated, etomoxir (20 μM) or BPTES (3 μM) was added to the plate 20 min prior to basal respiration measurements.

### Pyruvate-driven or glutamate-driven oxygen consumption rate (OCR) measurements in isolated mitochondria

A Seahorse Bioscience XF-96 extracellular flux analyzer was used to monitor mitochondrial oxygen consumption similar as previously described^56^. 5 μg of isolated skeletal muscle mitochondria suspended in a buffer containing 70 mM Sucrose, 220 mM D-mannitol, 10 mM KH_2_PO_4_, 5 mM MgCl_2_, 5 mM HEPES pH 7.2, 1 mM EGTA, and 0.2% fatty acid free BSA were attached to V3-PET seahorse plates by centrifugation at 2000 g for 20 min. Substrates of interest were at final concentrations of 10 mM Pyruvate/1 mM Malate or 10 mM Glutamate/1 mM Malate. A three-injection protocol was utilized with three replicate measurements taken between each injection. Each replicate consisted of a 1 min mix step, a 1 min equilibration step, and a 3 min measurement step. After basal measurements were acquired, maximum oxygen consumption was stimulated by the addition of 4 mM ADP and 1 μM FCCP. Mitochondrial pyruvate carrier (MPC) specific activity was inhibited by the addition of 10 μM UK5099 (MPC inhibitor^57^). Finally, Complex I activity was inhibited by 5 μM Rotenone and 2μM antimycin A. Oxygen consumption was normalized to mitochondrial protein loading.

### Metabolite Extraction and GC/MS Analysis

At the conclusion of a tracer experiment, the tracer media was removed from the culture wells, the cells were washed with a saline solution, and the bottom of the well was covered with cold methanol to lyse the cells and halt metabolism. Water containing norvaline at 5 mg/ml was charged to each well at a volume ratio of 1:2.5 relative to the methanol. The bottom of each well was scraped with a 1,000 ml pipette tip, and the cells were collected in 1.5 ml tubes. Cold chloroform containing 2 μg/ml of heptadodecnaoate was added to each sample at a 1:1 volume ratio relative to the methanol. The mixtures were vortexed, and the polar and nonpolar layers were separated and evaporated after centrifugation. Derivatized metabolites were analyzed using a DB-35MS column in an Agilent 7890A gas chromatograph coupled to a 5975C mass spectrometer. Details can be found in the Supplemental Experimental Procedures.

## Funding

This study was supported by grants from the National Natural Science Foundation of China (30900611 to C.W.) and supported by Young Talent Fund of University Association for Science and Technology in Shaanxi, China (20170404 to N.L.).

## Conflict of Interest

We don’t have any conflict of interest to declare.

## Notes

### Competing Interest Statement

The authors have declared no competing interest.

